# Novel Polymycoviruses Are Encapsidated in Filamentous Virions

**DOI:** 10.1101/2024.03.08.584110

**Authors:** Zhenhao Han, Jingjing Jiang, Wenxing Xu

## Abstract

*Polymycoviridae* is a new viral family that was established nearly five years ago, but their viral morphologies (naked or encapsidated) remain controversy since only one member namely, Colletotrichum camelliae filamentous virus 1 (CcFV1), was identified as being encapsidated in filamentous virions. Here, three novel dsRNA viruses belonging to the family *Polymycoviridae* were identified in three phytopathogenic fungal strains and tentatively named Pseudopestalotiopsis camelliae-sinensis polymycovirus 1 (PscPmV1), and Phyllosticta capitalensis polymycovirus 1 (PhcPmV1) and -2 (PhcPmV2), respectively. PscPmV1 and PhcPmVs have five or six genomic dsRNAs, ranging from 1055 to 2405 bp, encoding five or seven putative open reading frames (ORFs), of which ORF1 encodes an RNA-dependent RNA polymerase, ORF5 encodes a P-A-S-rich protein behaving as coat protein (CP); and dsRNAs 4 and 6 encode putative proteins with unknown functions and share no detectable identities with known viral sequences. Upon examination under transmission electron microscopy after purification from fungal mycelia, PscPmV1 and PhcPmVs were found to be encapsidated in filamentous particles, as was a known polymycovirus, Botryosphaeria dothidea RNA virus 1 (BdRV1), which was previously assumed to likely have no conventional virions. The morphology of PscPmV1 was further supported by the observation that its particles could be decorated by polyclonal antibodies against its CP and bound by immuno-gold particles conjugated to the specific CP antibody. Together with CcFV1 and BdRV1, PcsPmV1 PhcPmVs provide strong evidence to support the notion that polymycoviruses are encapsidated in filamentous virions constituted by P-A-S-rich CPs. Moreover, their biological effects on their fungal hosts were assessed.

**Significance statement:** *Polymycoviridae*, a recently established viral family, has raised questions about encapsidation. Here, we identify and characterize three novel polymycoviral dsRNA viruses in phytopathogenic fungal strains, tentatively named Pseudopestalotiopsis camelliae-sinensis polymycovirus 1, and Phyllosticta capitalensis polymycovirus 1 and -2, respectively. These polymycoviruses possess five or six genomic dsRNAs, ranging from 1055 to 2405 bp, with two encoding putative proteins of unknown functions and sharing no detectable identities with known viral sequences. Their morphologies indicate filamentous virions constituted by P-A-S-rich coat proteins, observed using immunosorbent electron microscopy combined with immune-gold labeling techniques. Additionally, Botryosphaeria dothidea RNA virus 1, previously assumed to lack conventional virions, is also shown to be encapsidated in filamentous particles. This study provides new evidence supporting the encapsidation of polymycoviruses into elongated and flexuous virions, significantly contributing to our understanding of the evolutionary particle architecture within the virosphere and expanding our knowledge of viral diversity and evolution.

## Introduction

The morphotypical diversity of viruses is an important trait that reflects their origin, taxon, evolution and host in the expanding virosphere considering that their morphotypical peculiarities have been influenced by the environment and the specific nature of the host (1–3). Filamentous virions have been widely observed in positive single-stranded RNA [(+)ssRNA)] virus families, such as *Closteroviridae*, *Potyviridae*, *Alphaflexiviridae*, *Betaflexiviridae*, and *Gammaflexiviridae*, as well as in some double-stranded (ds) DNA virus families like *Lipothrixviridae*, positive single-stranded (+ss) DNA virus families (e.g., *Inoviridae*), and negative single-stranded (-ss) RNA virus families (e.g., *Filoviridae*). However, filamentous virions are far less observed in dsRNA virus groups, with the exception of Colletotrichum camelliae filamentous virus 1 (CcFV1, also known as Colletotrichum camelliae polymycovirus 1) infecting a phytopathogenic fungus, even among those isolated from other organisms including protozoa and animals (4). It is striking that such a filamentous architecture has far less been observed in dsRNA virus groups, thus leaving the evolutionary relationships between the morphologies of (+)ssRNA and dsRNA viruses enigmatic since previous studies have revealed that dsRNA viruses may have repeatedly originated from distinct supergroups of (+)RNA viruses (5, 6).

The mycoviral family *Polymycoviridae* was established nearly five years ago, and their members are phylogenetically linked to both dsRNA and (+)ssRNA viruses, suggesting a potential origin from a (+)ssRNA viral ancestor belonging to clades three or six of the picorna-like superfamily (4, 7). Recently, the number of members belonging this family has significantly increased due to advancements in next-generation sequencing (NGS) technology. There members infect fungi (ascomycetes and basidiomycetes) and oomycetes (2, 4), indicating a dynamic genomic organization in terms of segment number (ranging from four to eight) and sequence (4, 5). Known polymycoviruses possess multipartite genomes ranging in size from 7.5 to 12.5 kbp, each containing one or two open reading frames (ORFs) (2, 4). They encode four conserved proteins, including an RNA-dependent RNA polymerase (RdRp), a hypothetical protein of unknown function (containing a conserved N-terminus and a zinc-finger motif), a putative methyltransferase (Met), and a proline-alanine-serine rich protein (PASrp) (2, 3, 7). Interestingly, the members Aspergillus fumigatus tetramycovirus 1 (AfuTmV1) from the human pathogenic fungus and Beauveria bassiana polymycovirus 1 (BbPmV1) from an entomopathogenic fungus seem to form non-conventional virions associated with PASrp in a colloidal form, as observed using atomic force microscopy (AFM) (5, 7); Penicillium digitatum polymycovirus 1 (PdPmV1) from a phytopathogenic fungus infecting citrus showed no detectable virus-like particles (8), while Botryosphaeria dothidea RNA virus 1 (BdRV1; also known Botryosphaeria dothidea polymycovirus 1, BdPmV1) from a phytopathogenic fungus infecting pear was associated with some short bacilliform virus-like particles and likely lacked non-conventional virions (9) and only CcFV1 is believed to have a filamentous capsid (4). Given the homology of PASrps among these polymycoviruses, it is conceivable to anticipate that homologous PASrps may lead to a similar virus morphology (10), prompting a controversy regarding whether the viral morphologies of this family are naked or encapsidated, and further morphological characterization of polymycoviruses is warranted (10).

Here, we present the isolation and characterization of three novel dsRNA viruses belonging to the family *Polymycoviridae*. These viruses were isolated from three strains of two different phytopathogenic fungi, infecting pear (*Pyrus communis* L) and tea [*Camellia sinensis* (L.) O.Kuntze] in China, respectively. Our findings indicate that they are encapsidated in filamentous virions constituted by PASrps, as observed using immunosorbent electron microscopy (ISEM) in combination with immune-gold labeling (IGL) techniques. This study offers new evidence supporting the encapsidation of polymycoviruses into elongated and flexuous virions.

## Results

### Six dsRNAs in *Pseudopestalotiopsis camelliae-sinensis* strain CYG1-2 compose the genome of a novel polymycovirus

Nucleic acids extracted from the mycelia of *Ps. camelliae-sinensis* strain CYG1-2 and CJP3-4 (Fig. 1A) were digested by S1 nuclease and fractionated on an agarose gel, revealing several dsRNAs in CYG1-2 but not in CJP3-4 (Fig. 1B). The dsRNAs were further analyzed on a PAGE gel to determine their numbers and sizes, revealing six bands (termed dsRNAs 1 to 6 according to their decreased sizes) in size ranging from 1.0 to 2.5 kbp, compared to the genomic RNA sizes of Colletotrichum fructicola RNA virus 1 (CfRV1) (Fig. 1C).

**Fig. 1.**
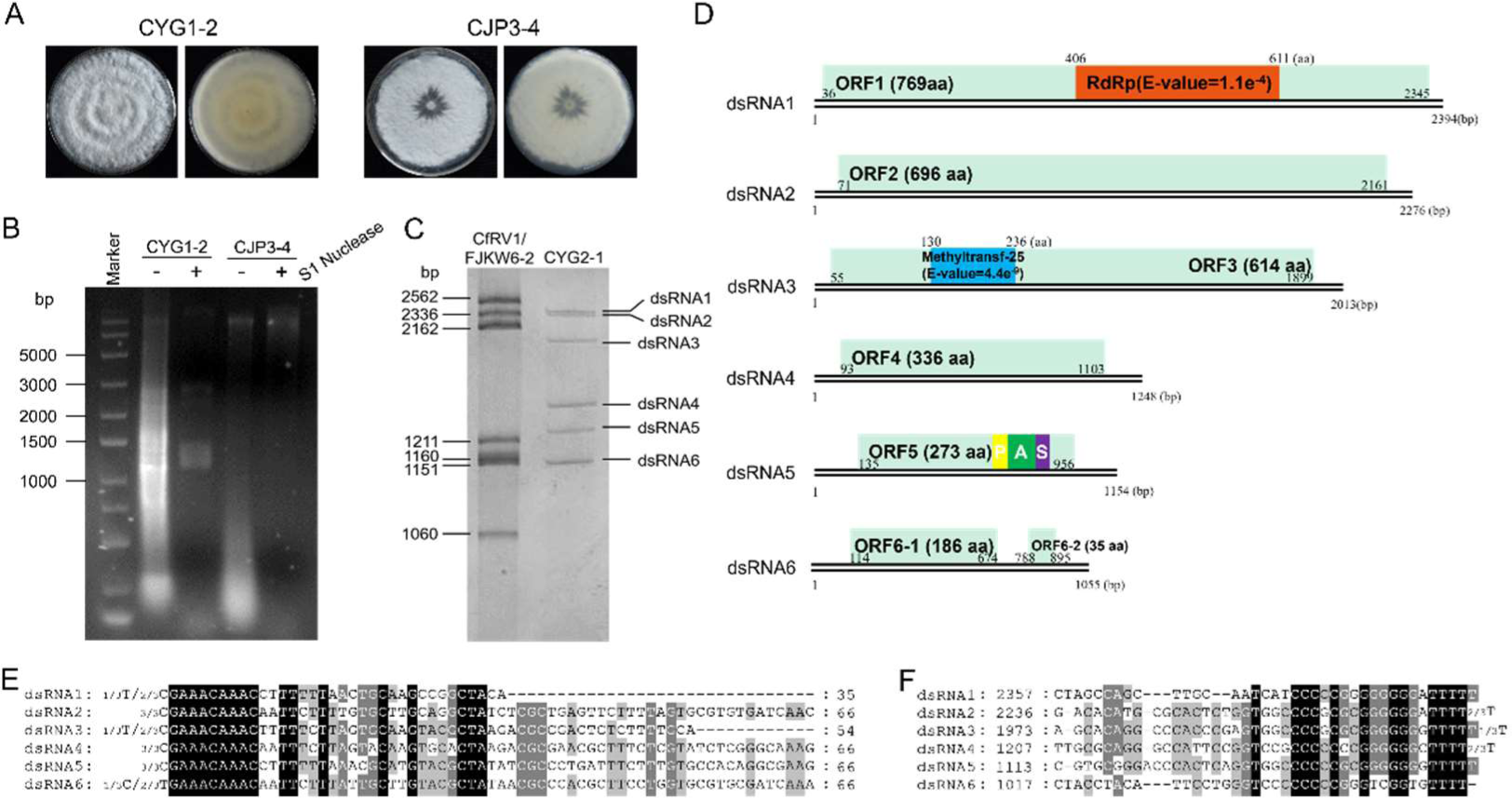
Fungal morphologies, nucleic acid electrophoresis, genomic characteristics, and multiple alignments of the terminal regions of dsRNAs 1-6 of Pseudopestalotiopsis camelliae-sinensis polymycovirus 1 (PcsPmV1). (A) Colonies of *Pseudopestalotiopsis camelliae-sinensis* strains CYG1-2 and CJP3-4 grown on PDA medium for 7 days. (B) Electrophoresis analysis of nucleic acids extracted from strains CYG1-2 and CJP3-4 without treatment (lane 2 and 4) and treated with S1 nuclease (3 and 5) on 1.5% agarose gel. (C) Electrophoresis analysis of nucleic acids extracted from strain CYG1-2 treated with S1 nuclease (right lane) on 6% polyacrylamide gel and Colletotrichum fructicola RNA virus 1 (CfRV1) belonging to *Hadakaviridae* as dsRNA marker (left lane). (D) Genomic organization of dsRNAs 1-6 showing putative open reading frames (ORFs) and untranslated regions (UTRs). (E and F) Conserved sequences of the 5’- and 3’-termini of the dsRNAs, respectively. Black, grey, and light grey backgrounds denote nucleotide identity of no less than 100%, 80% and 60%, respectively. The terminal nucleotides, along with their frequencies in the RACE experiments, are indicated adjacent to the strand ends.

The full-length cDNA sequences of dsRNAs 1 to 6 were determined by assembling partial cDNAs amplified from separately purified dsRNAs using RT-PCR with tagged random primers and RACE, revelaing 2394, 2276, 2013, 1248, 1154, and 1055 bp, respectively (Fig. 1D). Each dsRNA contains one (for dsRNAs 1 to 5) or two (dsRNA 6) putative ORFs (termed ORFs 1 to 6 including ORFs 6-1 and 6-2 correspondingly) on one of the strands, encoding seven putative proteins (termed P1 to P6 including P6-1 and -2 correspondingly) (Fig. 1D). BLASTp searches of P1 to P6 revealed that P1 to P3 and P5 share amino acid sequence identities of 47.97% (100% coverage, *E*-value = 0) to 58.93% (99% coverage, *E*-value = 0) with the RNA-dependent RNA polymerase (RdRp), hypothetical protein, methyl transferase (Met), and PAS-rich protein of Phaeoacremonium minimum tetramycovirus 1 (PmTmV1) and Metarhizium brunneum polymycovirus 1 (MbPV1) (Table S2), respectively . Whereas the remaining proteins (namely P4 and P6-1 and -2) had no detectable identities with viral proteins.

The 5’-untranslated regions (5’-UTRs) of the coding strands of dsRNA1 to dsRNA6 were 35, 70, 43, 92, 134 and 113 nt long (Fig. 1D), respectively, and shared 15% to 49% identity with each other, while the corresponding 3’-UTRs were 49, 115, 114, 145, 198 and 160 nt long (Fig. 1D), respectively, and shared 19% to 62% identity with each other. Both termini of dsRNAs 1 to 6 contain conserved sequences, including the first 5’-terminal nucleotides (GAAACAAAC) and the last 3’-terminal nucleotides (TTTT) (Fig.1E, F). Since dsRNAs 1 to 6 share conserved sequences at both termini, characteristic of segmented genomic components of a polymycovirus (2), dsRNAs 1 to 6 are determined as the genomic components of a novel polymycovirus, tentatively named Pseudopestalotiopsis camelliae-sinensis polymycovirus 1 (PcsPmV1). The genomic sequences have been deposited in GenBank under accession numbers PP359405 - PP3594010.

Putative protein functions for the PcsPmV1 ORFs were also inferred by a homology search using the Pfam database, with the results obtained agreeing with those from BLASTp searches (Table S2). PcsPmV1 P1 contains an RdRp domain belonging to the *RdRp_1* family. Sequence alignment allowed the identification of three amino acid motifs (IV, V and VI) also observed in AfuTmV1 and CcFV1, further supporting that P1 is an RdRp, with all of them containing the GDN triplet followed by Q rather than G/ADD in motif VI (which is normally invariant for (+) ssRNA viruses, Fig. S1A). No conserved domain was predicted in PcsPmV1 P2; meanwhile, alignment of P2 with other identified polymycovirus found a ZinC finger domain, which was rich in cysteine (C) (Fig. S1B). PcsPmV1 P3 contains a Met domain, functioning as an S-adenosyl methionine-dependent Met capping enzyme as predicted previously in AfuTmV1 and CcFV1, which is indicated by a principal lysine residue (K, Fig. S1C). No function could be tentatively ascribed to the remaining PcsPmV1 putative proteins due to a lack of reliable conserved motifs.Nonetheless, PcsPmV1 P5 has a high proportion of P (7.7%), A (13.9%), and S (8.1%) residues, resembling the PASrps encoded by AfuTmV-1 (P 6.8%, A 8.0%, S 9.7%), BdRV1 (P 3.2%, A 9.7%, S 7.9%), BbPmV-1 (P 6.2%, A 13.5%, S 7.6%) and CcFV1 (P 7.6%, A 13.4%, S 10%), which is a putative CP.

### Five dsRNAs in *Phyllosticta capitalensis* strains compose the genome of two novel polymycoviruses

Five dsRNAs (termed dsRNAs 1 to 5) were detected in *Ph. capitalensis* strain DHP2-1 (Fig. 2A) and their lengths were determined as described for PcsPmV1 (Fig. 2B), which are 2344, 2187, 1936, 1443 and 1223 bp, respectively (Fig. 2C). Each of the five dsRNAs contains five putative ORF (termed ORFs 1 to 5), encoding five putative proteins, designated as P1 to P5, respectively (Fig. 2C). BLASTp searches of P1 to P5 revealed that P1 to P3 share amino acid sequence identities of 55.36% (96% coverage, E-value = 0), 42.78% (99% coverage, E-value = 0), and 44.21% (94% coverage, E-value = 2e-168) with RdRp, hypothetical protein, and Met of AfuPmV1, respectively; P4 shares no detectable identities with viral proteins. while P5 shares an amino acid sequence identity of 42.48% (96% coverage, E-value = 8e-62) with the PAS-rich protein of Erysiphe necator associated polymycovirus 4 (EnaPmV4), supporting its classification as a putative CP.

**Fig. 2.**
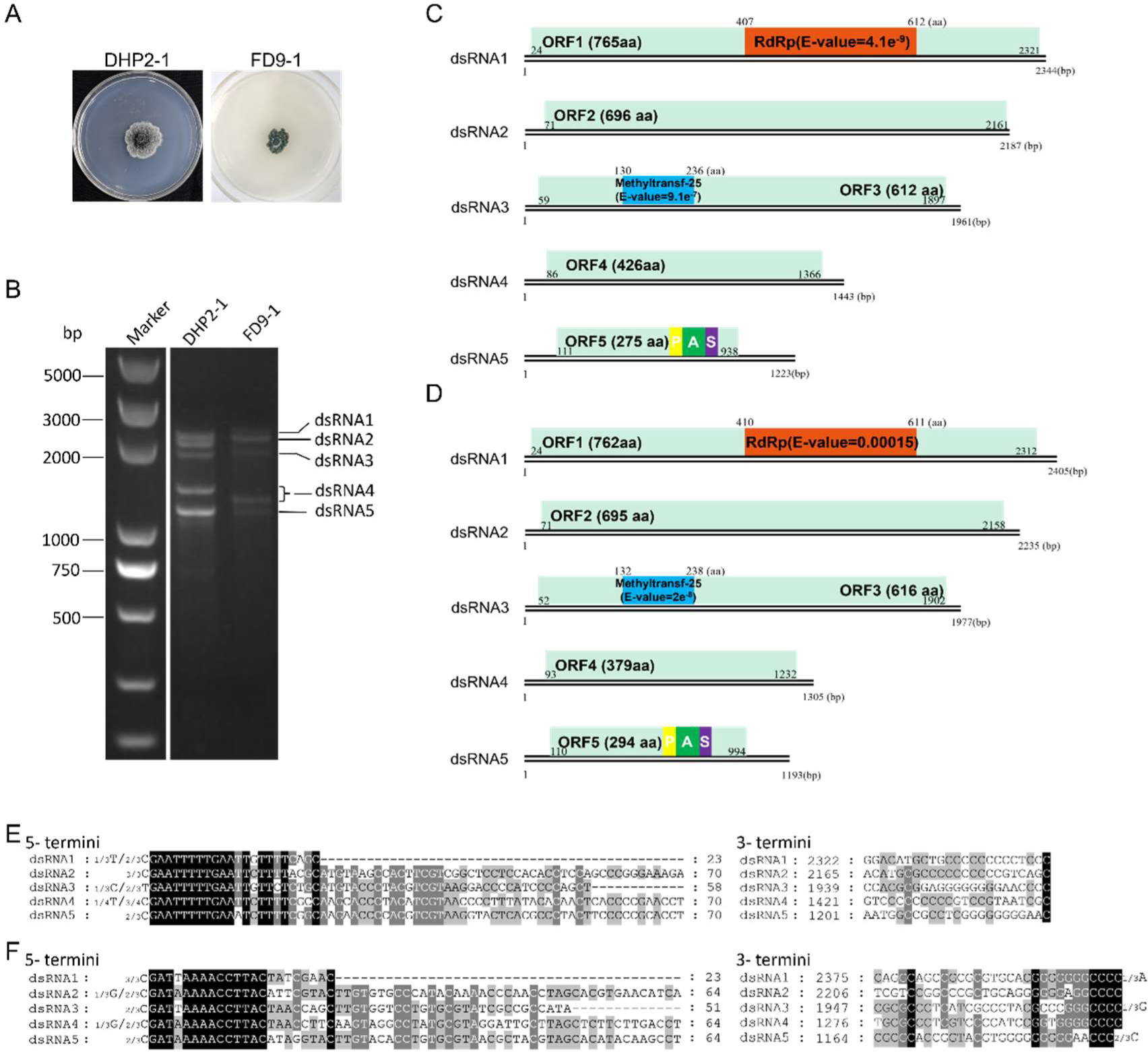
Fungal morphologies, genomic characteristics, and multiple alignments of the terminal regions of dsRNAs 1-5 of Phyllosticta capitalensis polymycovirus 1 (PhcPmV1) and -2. (A) Colonies of *Ph. capitalensis* strains DHP2-1 and FD9-1 grown on PDA medium for 7 days. (B) Electrophoresis analysis of nucleic acids extracted from strains DHP2-1 and FD9-1 on 1.0% agarose gel. (C and D) Genomic organization of dsRNAs 1-5 of PhcPmV1 and -2, respectively. (E and F) Conserved sequences of the 5’- and 3’-termini of the dsRNAs of PhcPmV1 and -2, respectively.

Both termini of dsRNAs 1 to 5 contain conserved sequences, including the first 5’-terminal nucleotides (GAATTTTTGAA) and the last 3’-terminal nucleotide (C) (Fig. 2E). Similar to PcsPmV1, dsRNAs 1 to 5 are determined as the genomic components of a novel polymycovirus, tentatively named Phyllosticta capitalensis polymycovirus 1 (PhcPmV1). The genomic sequences have been deposited in GenBank under accession numbers PP359411 - PP359415.

In *Ph. capitalensis* strain FD9-1, five dsRNAs (termed dsRNAs 1 to 5) were detected and determined to be of sizes 2405, 2235, 1977, 1305, and 1193 bp, respectively (Fig. 2D). These dsRNAs contain five putative ORF (termed ORFs 1 to 5), each on one strand, encoding five putative proteins (P1 to P5 correspondingly) (Fig. 2D). BLASTp searches of P1 to P5 revealed that P1 to P3 share amino acid sequence identities of 55.72% (98% coverage, E-value = 0) , 40.27% (94% coverage, E-value = 2e-114), and 47% (96% coverage, E-value = 2e-173) with RdRp, hypothetical protein and Met of AfuPmV1, respectively; P4 shares no detectable identities with viral proteins, while P5 shares an amino acid sequence identity of 44.77% (92% coverage, E-value = 2e-58) with the PAS-rich protein of EnaPmV4, indicating that it is a putative CP. Similarly, dsRNAs 1 to 5, containing conserved 5’-(GATA/TAAAACCTTAC) and 3’-terminal nucleotides (CCC) (Fig. 2F), are determined to be the genomic components of a novel polymycovirus, tentatively named Phyllosticta capitalensis polymycovirus 2 (PhcPmV2). The genomic sequences have been deposited in GenBank under accession numbers PP3594116-PP359411.

Putative protein functions for the ORFs of PhcPmVs (including both PhcPmV1 and -2) were inferred through a homology search using the Pfam database, which corroborated results obtained from BLASTp searches. Each PhcPmV P1 contains an RdRp domain belonging to the RdRp_1 family, similar to AfuTmV1 and CcFV1. However, no conserved domains were predicted in PhcPmV P2. Each PhcPmV P3 contains a Met domain. No specific function could be tentatively ascribed to the remaining putative proteins of PhcPmVs due to a lack of reliable conserved motifs. Nonetheless, PhcPmV P5 has a high proportion of P (9.5% for PhcPmV1 and 7.1% for PhcPmV2), A (10.5% and 10.5%), and S (7.6% and 4.8%) residues, resembling the PASrps encoded by AfuTmV-1, BdRV1, BbPmV-1, and CcFV1.

### PcsPmV1 and PhcPmVs are phylogenetically related to polymycoviruses assumed to be naked

Phylogenetic analyses of the putative RdRps and CPs of PcsPmV1 and PhcPmVs, along with the corresponding proteins of other polymycoviruses revealed that they are all new members belonging to the family *Polymycoviridae*. Based on the RdRp phylogenetic topology, PcsPmV1 and PhcPmV1 are closest phylogenetically to BbPmV1 and MoPmV1, while PhcPmV2 is closest to BdRV1, AfuPmV1, and AspTMV1; whereas based on the CP phylogenetic topology, PcsPmV1 and PhcPmV1 are closest to MoPmV1 and BdRV1, while PhcPmV2 closest to AfuPmV1 and AspTMV1 (Fig. 3A, B). The results indicate that PcsPmV1 and PhcPmVs are phylogenetically related to the polymycoviruses AfuPmV1 and BdRV1, which were assumed to be naked.

**Fig. 3.**
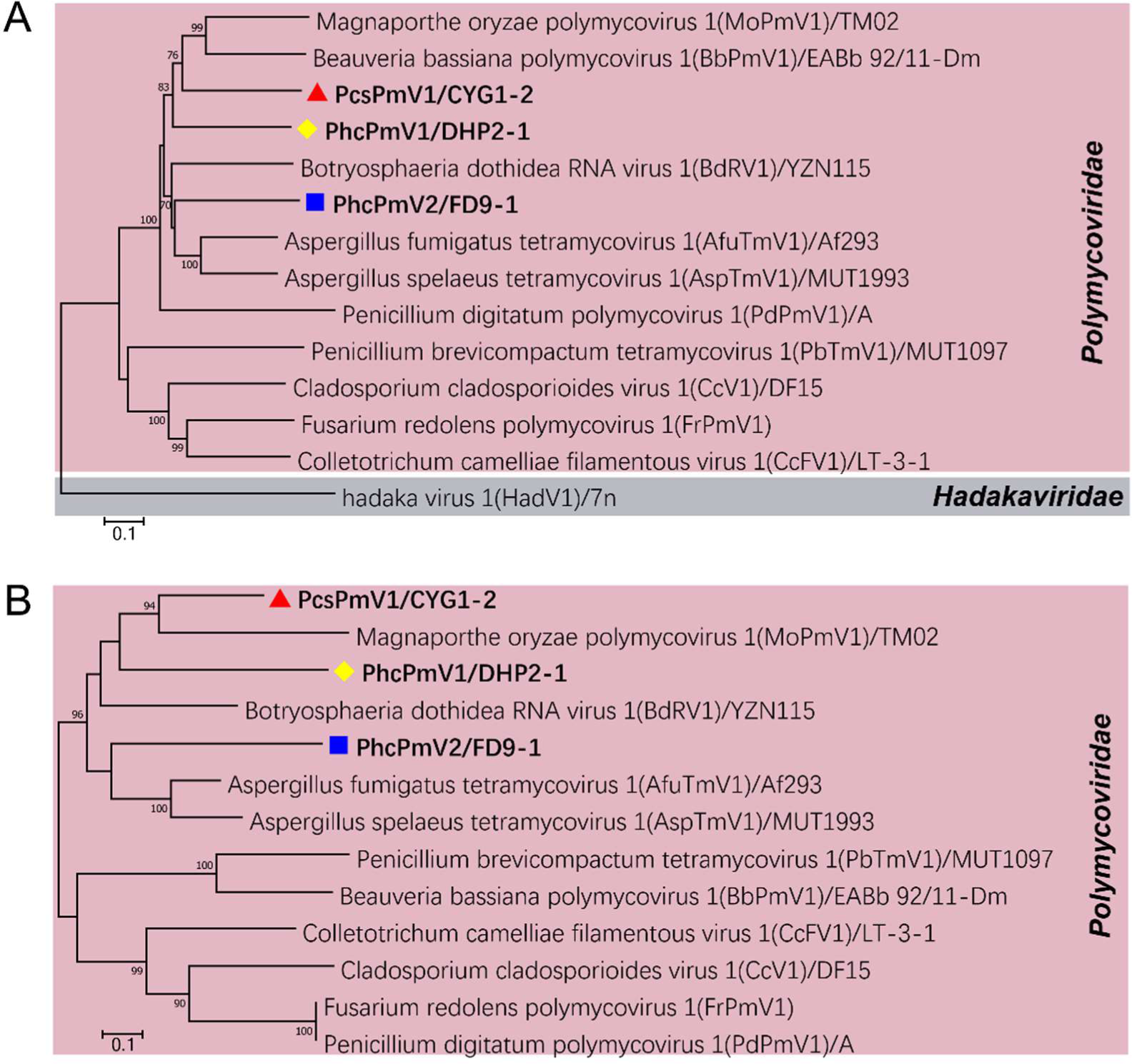
Phylogenetic analysis of Pseudopestalotiopsis camelliae-sinensis polymycovirus 1 (PcsPmV1) and PhcPmVs, respectively. (A and B) ML phylogenetic trees constructed based on the deduced the RNA-dependent RNA polymerase (RdRp) sequences (A) and capsid protein (CP) sequences (B) of PcsPmV1, PhcPmVs, and other polymycoviruses.

### PcsPmV1 and PhcPmV1 are associated with filamentous virus-like particles

To ascertain whether these polymycoviruses are encapsidated, PcsPmV1 and PhcPmV1 were chosen for viral particle determination based on the clustering of their CPs with BdRV1 and AfuPmV1respectively, which are assumed to be naked. Viral preparation of PcsPmV1 was purified from the mycelia of *Ps. camelliae-sinensis* strain CYG1-2 by ultracentrifugation in stepwise sucrose gradients (60–10% with 10% sucrose increments) and examined by transmission electron microscopy (TEM). The TEM analysis revealed filamentous virus-like particles longer than 200 nm, being predominantly found in the 30% and 40% sucrose fractions (Fig. 4A, B). Considering that the observed short particles were most likely fragments of the longer ones, only those longer than 400 nm were considered. Measurements of 11 filamentous virus-like particles above that limit showed widths ranging from 16.4 (number 8) to 20.3 (number 3) nm (Fig. 4C) and lengths from 412.1 (number 4) to 3307.1 nm (number 11, Fig. 4C). Similarly, filamentous particles were observed under TEM for the viral preparation of PhcPmV1 from the mycelia of *Ph. capitalensis* strain DHP2-1, predominantly found in the 20% sucrose fractions. These particles had widths ranging from 12.72 to 14.88 nm and lengths from 666.36 to 1912.69 nm (Fig. S3).

**Fig. 4.**
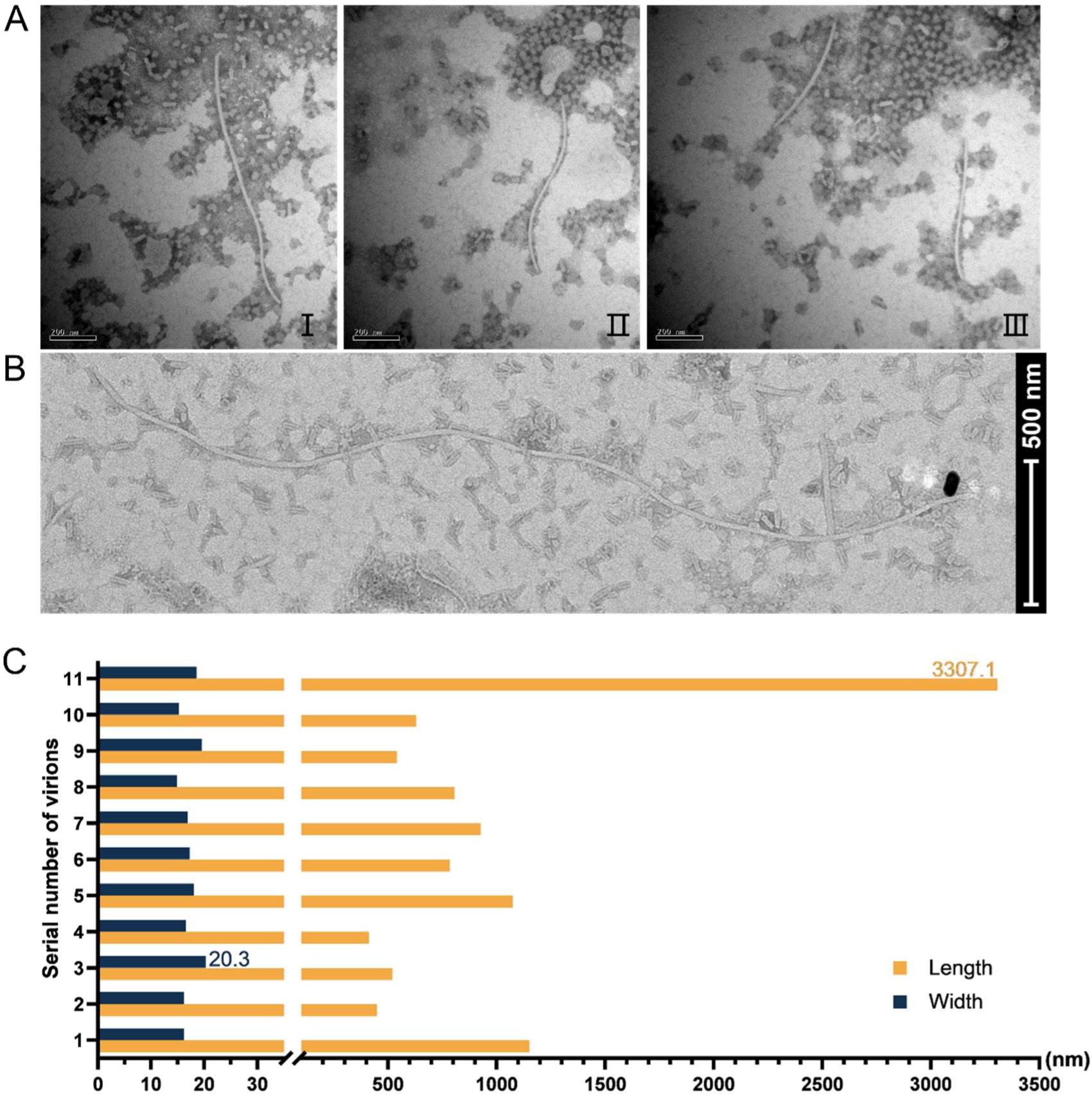
Representative transmission electron microscopy (TEM) images of virus-like particles extracted from *Ps. camelliae-sinensis* strain CYG1-2 and a histogram of the sizes of particles longer than 400.0 nm. (A) Representative virus-like particles extracted from strain CYG1-2 corresponding to the 30% fraction following sucrose gradient centrifugation. The arrows indicate empty particles that might have lost dsRNA. Scale bars, 200 mm. (B) Virus-like particles measuring 3307.1 nm in length and 18.6 nm in width observed in the 40% sucrose fraction. Scale bar, 500 nm. (C) A histogram depicting the sizes of particles longer than 400.0 nm from strain CYG1-2 in fractions corresponding to 30% and 40% sucrose fractions. The numbers on the vertical axis represent counts of virus-like particles.

### ORF5-coding proteins and the dsRNAs compose the virus-like particles of PcsPmV1

Agarose gel electrophoresis of the nucleic acids extracted from the 10–60% sucrose gradient fractions (at 10% sucrose increments) showed that the typical pattern of PcsPmV1 dsRNAs 1–6 were mostly recovered from the 40% fractions (Fig. 5A), which suggests that dsRNAs 1–6 are all encapsidated in the virus particles of PcsPmV1. No dsRNA bands were recovered from any of the gradient fractions from mycelial extracts of strain CJP3-4 performed in parallel (Fig. 5A).

**Fig. 5.**
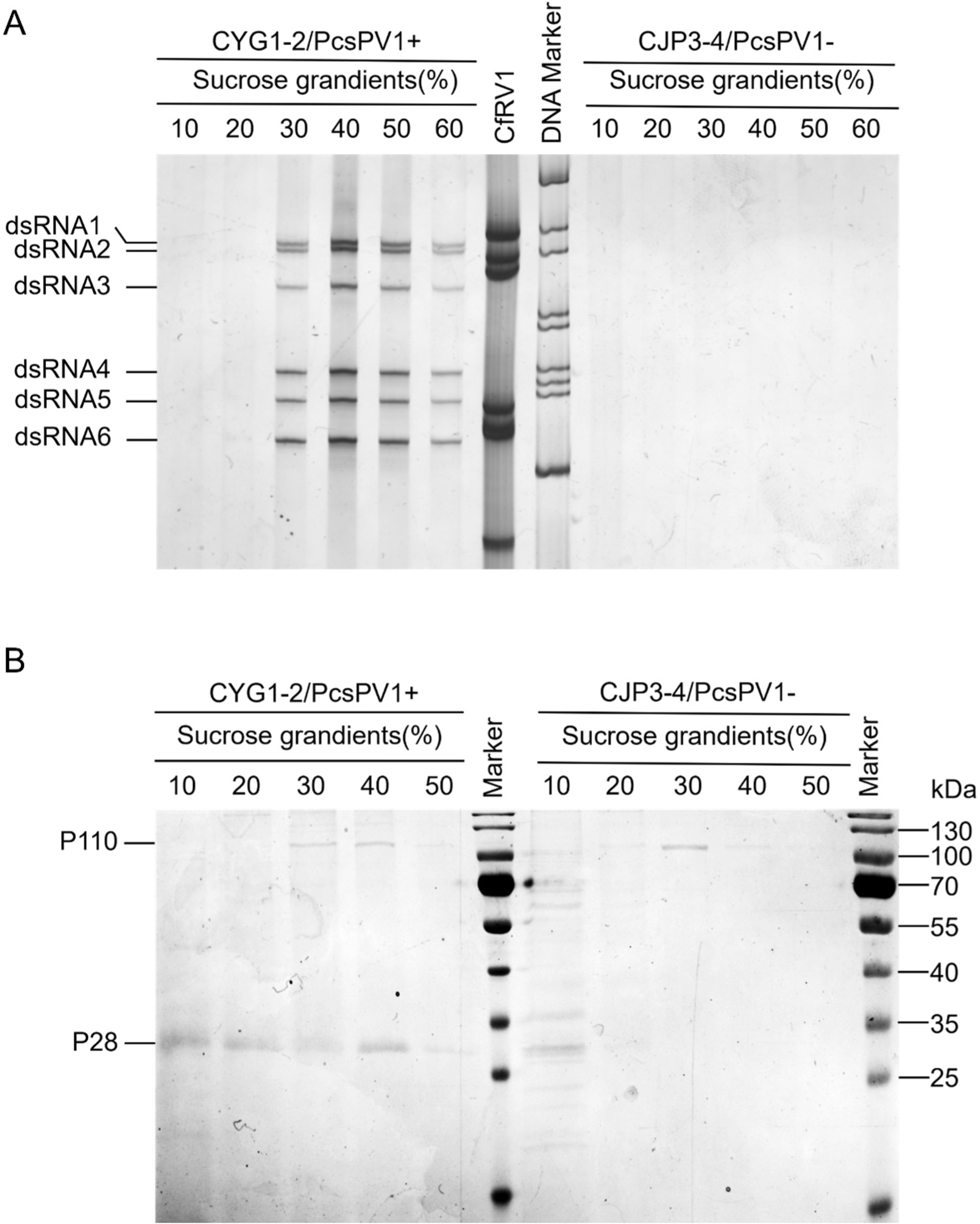
Analysis of nucleic acids and proteins associated with virus-like particles. (A) Polyacrylamide gel electrophoresis (PAGE) analysis of dsRNAs extracted from purified virus-like particles from 10% to 60% sucrose fractions at 10% increments from strain CYG1-2 (lane 1-6) and CJP3-4 (lane 9-14). (B) SDS-PAGE analysis of proteins extracted from 10% to 50% sucrose gradient fractions (with 10% increments) of strains CYG1-2 and CJP3-4.

SDS-PAGE analysis of the proteins from the 10-50% sucrose fractions of strain CYG1-2 revealed the presence of two dominant bands with estimated molecular masses of 110 and 28 kDa in the 30-50% fractions (Fig. 5B), tentatively termed p110 and p28, respectively. Of them, p110 instead of p28 was also observed in the control strain CJP3-4, suggesting p110 is a host protein (Fig. 5B). To further examine the nature of p28, the corresponding protein band in the 40% sucrose fraction was excised from the gel and subjected to peptide mass fingerprinting (PMF) analysis, which revealed that a total of 133 peptides generated from p28 matched P5 in 90% coverage (Table S3). This finding suggests that p28 is the structural protein P5 encoded by PcsPmV1 ORF5 (see below).

### PcsPmV1 P5 forms the capsid of virus particles

PcsPmV1 P5 was expressed and purified from *E.coli* BL21 (DE3), and injected into mice to trigger production of a polyclonal antibody (PAb-P5) against this protein. The antibody strongly and specifically recognized P5 from the 30% ∼ 50% sucrose fractions of strain CYG1-2, while no reactivity was observed with protein extracts from strain CJP3-4 (Fig. 6B). These results indicate that the purified P5 protein used as antigen was indeed uncontaminated with host proteins. Western blotting and indirect enzyme-linked immunosorbent assay (ELISA) analysis revealed a titer of ∼128,000-fold dilution for PAb-P5, with an optimum at 2000-fold dilution (Fig. 6A).

**Fig. 6.**
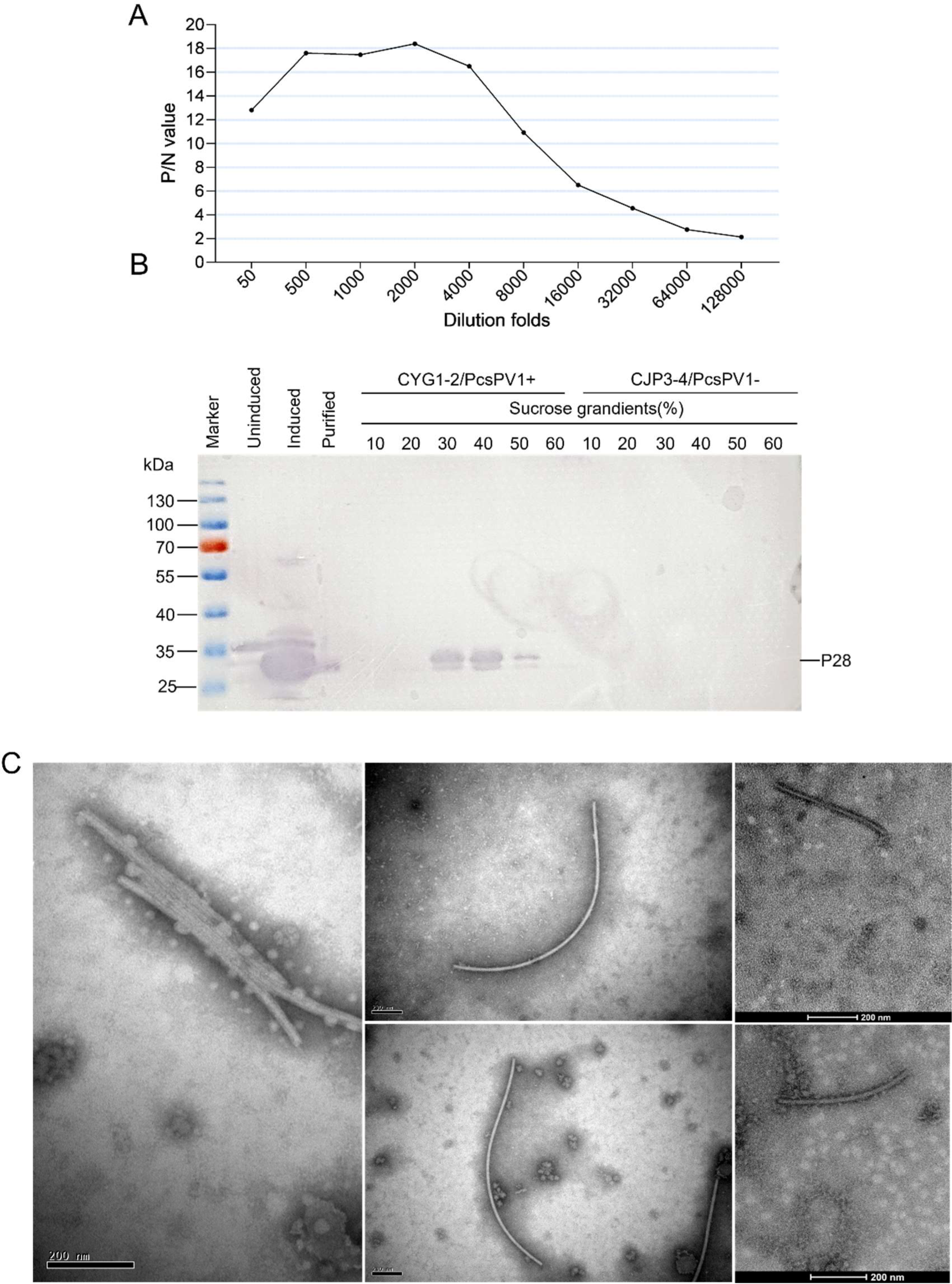
Titer quantification of a polyclonal antibody against PcsPmV1 P5, western blot analysis, and immunosorbent electron microscopy (ISEM) analysis of virus particles. (A) Titer quantification of PAb-P5 by indirect indirect enzyme-linked immunosorbent assay (ELISA) against the P5 (namely p28) protein from fractions after sucrose gradient centrifugation using PAb-P5 diluted from 50-to 128,000-fold. P/N represents the ratio of the absorbance values of the positive sample (PcsPmV1-infected CYG1-2) to the negative sample (virus-free CJP3-4) at a wavelength of 405 nm. (B) Western blot analysis of the proteins obtained from the *Escherichia coli* BL21 containing the reconstructed vector, pET-28a-P5-His (lane 2-4), and from strain CYG1-2 in 10-60% sucrose fractions (lane 5-10) after sucrose gradient centrifugation, and from strain CJP3-4 in the 10-60% sucrose fractions (lane 11-16) using the antibody against PcsPmV1 P5 (PAb-P5). ‘Uninduced’, ‘induced’, and ‘purified’ denote the proteins extracted from *E. coli* BL21 growing without an inducer, growing with an inducer, and after affinity purification, respectively. (C) ISEM analysis of virus-like particles derived from 30% and 40% fractions following sucrose gradient centrifugation of strain CYG1-2. The ISEM images show that the virus-like particles are decorated by PAb-P5 at a 4000-fold dilution. Scale bars, 200 mm.

To further confirm that the PcsPmV1 dsRNAs were encapsidated, the filamentous virus-like particles prepared by sucrose gradient centrifugation were subjected to immunosorbent electron microscopy (ISEM). The virus-like particles from the 40% sucrose fractions of strain CYG1-2 were all clearly decorated by PAb-P5 (Fig. 6C).

### Immuno-gold labeling confirms the capsid of PcsPmV1

To further demonstrate these virus particles were PcsPmV1, additional observation was done by immuno-gold labeling. These virus particles coated by PAb-P5 were clearly coated with some goat-anti-mouse gold-IgG (Fig. 7), which shows the virus rod had antigenic epitopes to which the gold-labeled antibodies could bind. Measurements of 10 decorated virus-like particles longer than 500 nm revealed widths ranging from 17.1 to 23.5 nm, matching the sizes observed by TEM. This result demonstrates the virus particles associated with CYG1-2 are encapsidated in the capsid encoded by P5 from PcsPmV1.

**Fig. 7.**
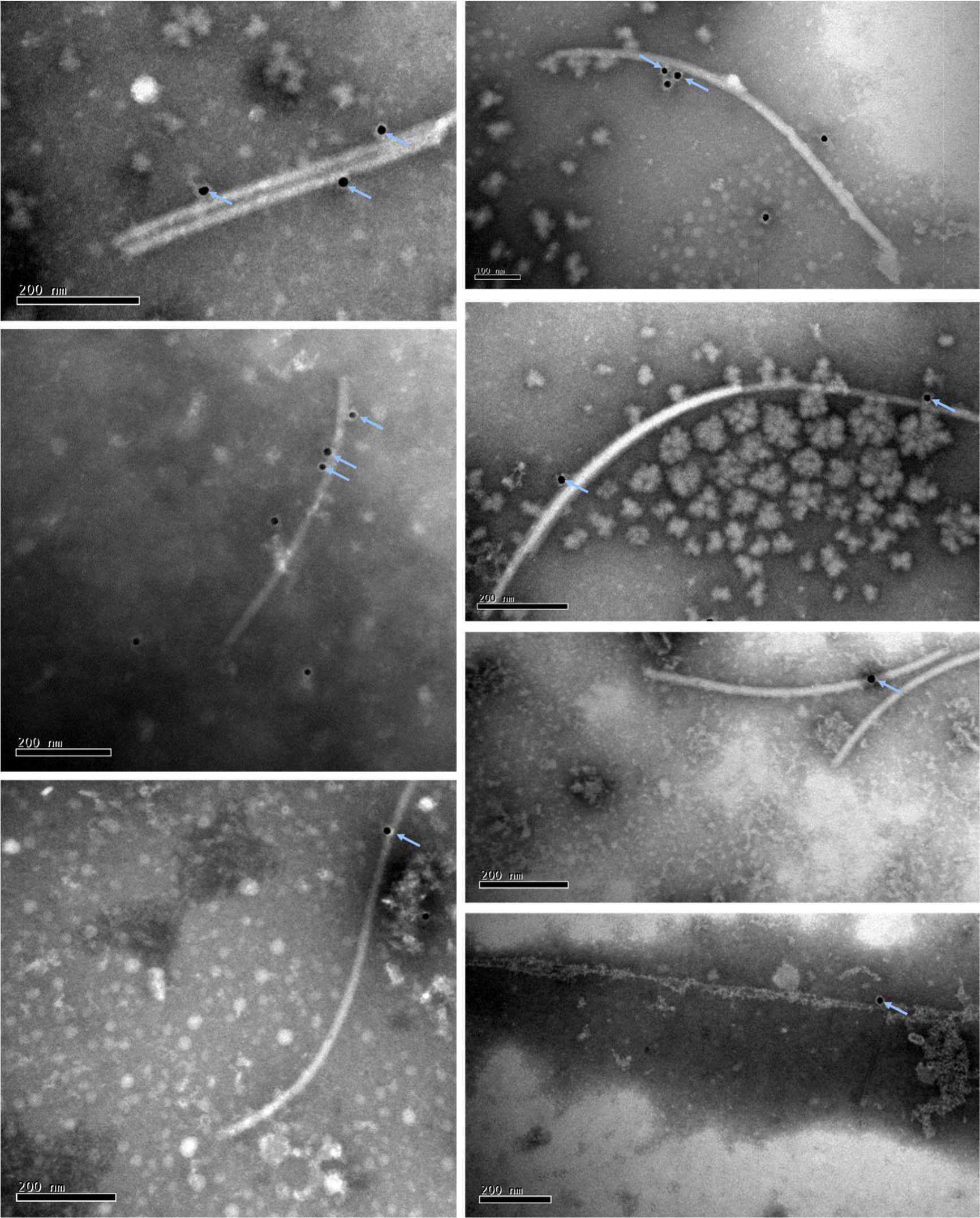
Immuno-gold labeling analysis of virus particles. Viruse particles labeled with PAb-P5 and a 15 nm goat anti-mouse IgG colloidal gold conjugate. The classic black dots (indicated by blue arrows) on the virions indicate that virus particles are positively labeled with immuno-gold.

### PcsPmV1 is related to stress response of the fungal host

Among 25 sub-isolates obtained by single conidium cultivation from PcsPmV1-infected strain CYG1-2, one PcsPmV1-cured isolate, nominated CYG1-2S9, was confirmed as being PcsPmV1-free by electrophoretic analysis of dsRNAs and RT-PCR amplification assay of a 632 bp fragment of PcsPmV1 dsRNA1 (Fig. S3A, B). The phenotypes of the PcsPmV1-infected CYG1-2 and PcsPmV1-free CYG1-2S9 together with their growth rates were observed and measured over a 6 d incubation period on PDA media (Fig. S5A, B). Apart from a minor reduction in the intensity of aerial mycelium in CYG1-2S9, there were no significant differences in morpholigies or growth rates between CYG1-2S9 and its virus-infected parent strain.

Additionally, to determine the role of PcsPmV1 in fungal stress responses, CYG1-2 and CYG1-2S9 were cultured on PDA amended with various cellular stress agents, including those involved in sugar metabolism (1 M sorbitol and 1 M glucose), cell wall disruption (0.01% SDS), osmotic stress (1 M KCl and 1.5 M NaCl), and the calcium pathway (0.5 M CaCl_2_). PcsPmV1-infected CYG1-2 showed stronger sensitivity to high concentrations of SDS and a slightly higher tolerance to high concentrations of KCl, CaCl_2_, sorbitol and glucose (Fig. S5C). These results showed that PcsPmV1 interfere with *Ps. camelliae-sinensis* and exert regulatory roles *in vivo* under different stress conditions.

### Filamentous virus-like particles are associated with BdRV1

To determine if BdRV1, containing five segmented genomic components dsRNAs 1 o 5, is encapsidated since it was previously considered without conventional virions (9) , the presumed virions were purified from mycelia of *Botryosphaeria dothidea* strain XA-3 by ultracentrifugation in a stepwise sucrose gradient (50–10% with 10% increments). Examination of the gradient fractions by TEM revealed filamentous virus-like particles with widths and lengths of 21–34 nm and 950–1500 nm, respectively (Fig. 8A and D), with the longest particle size being 1570×31 nm (Fig. 8B) and the shortest 950×21 nm (Fig. 8C).

**Fig. 8.**
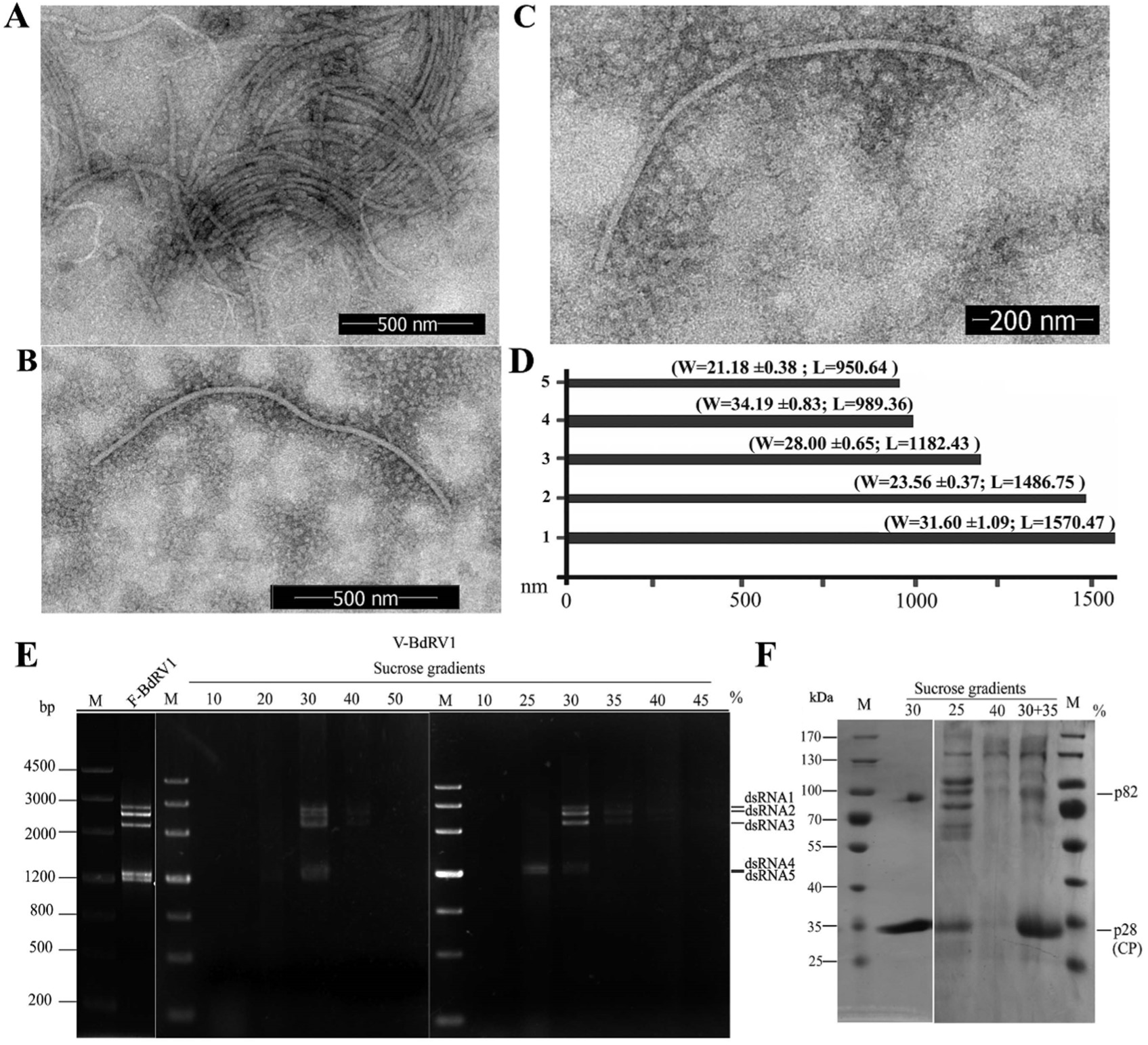
Virus-like particles, dsRNAs, and proteins of Botryosphaeria dothidea RNA virus 1 (BdRV1) extracted from the gradient fraction following sucrose gradient centrifugation. (A-C) Electron micrograph of virus-like particles purified from *Botryosphaeria dothidea* strain XA-3 from the gradient fraction corresponding to 30%, 25%, and 35% sucrose, respectively. (D) Histogram showing the sizes of the five longest particles. (E) Agarose gel electrophoresis analysis of the dsRNAs extracted from purified virus-like particles of BdRV1 from the gradient fraction ranging from 10% to 50% sucrose (V-BdRV1), and mycelia of strain XA-3 (F-BdRV1). ‘V-’ and ‘F-’ indicate dsRNAs extracted from virus-like particles and fungal mycelia, respectively. M, DNA size marker. (F) SDS-PAGE analysis of proteins extracted from purified particles of BdRV1 from the gradient fraction corresponding to 25% to 40% sucrose (30+35, a mixture of both sucrose layers). M, protein molecular weight marker.

Analyses by agarose or PAGE of the nucleic acids extracted from each gradient fraction revealed that the typical pattern of dsRNA1 to dsRNA5 previously isolated from XA-3 mycelium was predominantly recovered from the 30% sucrose fraction (Fig. 8E, left panel). A more resolute sucrose gradient centrifugation using 5% sucrose increments confirmed that the typical dsRNA1 to dsRNA5 profile was found in the 30% sucrose fraction and showed that the smallest and largest RNAs were predominantly found in the lower (25%) and higher (35%) sucrose fractions, respectively (Figs 8E, right panel). Virus-like particles from these fractions differed in size, with the largest (Fig. 8B) and smallest (Fig. 8C) particles recovered from 25% and 35% sucrose fractions, respectively. These data strongly suggest that dsRNA1 to dsRNA5 are separately encapsidated in elongated and flexuous virions, and their sizes depend on the length of the encapsidated dsRNAs.

SDS-PAGE protein analysis of the gradient fraction corresponding to 30% sucrose revealed the presence of two bands generated by proteins with estimated molecular masses of 28 (p28) and 82 kDa (p82) (Fig. 8F, left panel). Additional bands in size ranges of 70 to 170 KDa were observed when the gradient fractions corresponding to 25%, 35% and 40% of sucrose were analyzed, making the univocal identification of p82 difficult (Fig. 8F, right panel). However, p28 was always clearly discernible, and the signal intensity in each fraction (Fig. 8F) paralleled that of the dsRNA in the same fractions determined by agarose or PAGE analysis (Fig. 8E). To further examine the nature of these proteins, p82 and p28 (Fig. 8F, 30% line) were eluted from the gel and subjected to PMF analysis, in which a total of 37 and 15 peptide fragments were characterized, respectively.Among them, the peptide fragments from p82 matched proteins from fungi (most belonging to Ascomycota), with six peptides covering 10% of a protein from *Macrophomina phaseolina* (a fungus belonging to the family Botryosphaeriaceae), suggesting that p82 was a fungal protein (Table S4). The peptide fragments from p28 matched most (83.6%) of the amino acid sequence predicted to be encoded by dsRNA4 (Table S5). These data confirmed that p28 was indeed the structural proteins, namely CP. Taken together, these data strongly suggest that BdRV1 is encapsidated in flexuous and elongated virions, with ORF4 encoding its CP.

## Discussion

In this study, we identified three dsRNA virus clustered in the family *Polymycoviridae* and tentatively named them PcsPmV1, PhcPmV1, and PhcPmV2 which forms filamentous particles. Several experiments were conducted to confirm PcsPmV1 and PhcPmVs are indeed encapsidated in elongated and flexuous particles. (i) TEM of mycelial preparations from PcsPmV1- and PhcPmV-infected fungal strains revealed filamentous particles in the sucrose gradient fractions where the structural proteins and the genomic dsRNAs migrated, while these virus particles and structural proteins were absent in virus-free strain. (ii) The polyclonal antibody PAb-P5, raised against the structural protein P5, specifically recognized a structural protein of ∼28.6 kDa in specific sucrose gradient fractions, while exhibiting no reactivity with host proteins. Moreover, PAb-P5 decorated the filamentous particles in sucrose gradient fractions of PcsPmV1-infected strain. (iii) An immuno-gold labeling approach further confirmed that the filamentous particles could be recognized by PAb-P5. Taken together, these data provide strong evidence to support the encapsidation of PcsPmV1 and PhcPmVs dsRNAs in filamentous virions constituted by the CP formed by P5.

In previous studies, distinct virus forms have been reported for the members of *Polymycoviridae*. Among them, AfuPmV1 was reported to have non-conventional virions associated with PASrp in a colloidal form, BdRV1 associated with some short bacilliform virus-like particles, and CcFV1 filamentous (4, 5, 7–9). When their similarity was checked, their PASrps showed relatively high homology, ranging from 33% to 53%. It is conceivable to anticipate that these polymycoviruses should have a similar virus morphology, i.e., filamentous particles. We concluded that the short bacilliform virus-like associated with BdRV1 should be the broken fragments of the filamentous particles, which are easily observed in the filamentous virion purification process, e.g., for CcFV1 (4) and PcsPmV1 (Fig. 4B). PdRV1 virions failed to be detected ever through the authors tried to purify the related particles, and it could not be excluded the protocol limitation were utilized or the sample was selected in an unsuitable period for virion purification, since we have observed that the virion accumulation in high titer needs long periods for both CcFV1 and BdRV1. Here we characterize three novel polymycoviruses (PcsPmV1, and PhcPmV1 and -2) from different strains of two different phytopathogenic fungi (*Ps. camelliae-sinensis* and *Ph. capitalensis*) that were isolated from two different plants (pear and tea), which are cultivated in two different provinces (Hubei and Henan) at a far distance, but all of them form same filamentous morphologies even they had no related geography and host origins. Moreover, BdRV1, previously being considered without conventional virions (9), is discovered with filamentous particles. Together with CcFV1, we concluded that polymycoviruses are normally encapsidated in filamentous virions.

The putative P1 proteins of PcsPmV1 and PhcPmVs are predicted to be RdRps that additionally contain the GDNQ motif characteristic of the L genes of rhabdoviruses and paramyxoviruses within the order Mononegavirales (11). A similar feature was previously reported in the dsRNA viruses AfuTmV1 (7), BdRV1 (9), BbPmV1, BbPmV2 and CcFV1 (4), suggesting that, like the GDD motif in (+) ss RNA viruses, the RdRp GDNQ motif of PcsPmV1 and PhcPmVs could have important functions in metal ion coordination and nucleotide substrate binding during dsRNA replication (7). The putative protein P2 possesses a zinc finger-like motif, which has also been shown in P2 sequence of AfuTmV1 (7). The putative protein P3 is predicted to be an S-adenosyl methionine-dependent Met capping enzyme, as suggested for AfuTmV1 and CcFV1, and it might be involved in a cap-snatching mechanism (4, 7, 12). According to SDS-PAGE, PMF, ISEM and IGL analyses, the P-A-S-rich protein is the viral CP, constituting the viral particles. The functions of the remaining putative proteins are unclear due to a lack of sequence similarity with known proteins.

In previous studies, dsRNA viruses have been considered to have evolved from distinct supergroups of (+)RNA viruses, as their RdRps display features similar to those of different (+)RNA viruses (6, 13). In this context, the phylogenetic analysis of RdRps of (+)RNA and dsRNA viruses of eukaryotes indicates that polymycoviruses appear in an intermediate position between those of (+)ssRNA viruses (such as caliciviruses, picornaviruses, astroviruses, hypoviruses, and potyviruses) and dsRNA viruses, being closer to that of caliciviruses (4, 7). It is interesting to note that, in the evolutionary linkage, caliciviruses, picornaviruses and astroviruses from animals form isometric virions and are located in the leading position, followed by hypoviruses, which are naked from fungi, and lagged behind by filamentous potyviruses from plants (14, 15). However, the CPs of PcsPmV1, PhcPmVs, and other polymycoviruses show no detectable sequence similarity with proteins from known filamentous (+) ssRNA viruses that infect plants, including members of the families *Flexiviridae*, *Potyviridae* and *Closteroviridae*. This suggests that they should not have originated from the same single ancestral protein. Therefore, this study provides an important clue about the morphological origins and evolution of the family

*Polymycoviridae*, suggesting that they most likely originated from naked *Hypoviridae* viruses after a filamentous particle was evolutionarily obtained by an ancient hypovirus, similar to filamentous *Potyviridae* viruses, both of which were derived from *Astroviridae* after their star-like virions were shed.

In summary, the findings regarding PcsPmV1 and PhcPmVs strongly support the notion that polymycoviruses are encapsidated in filamentous virions. Importantly, this study sheds light on the morphological origins and evolution of the family *Polymycoviridae* and other dsRNA viruses. It represents a significant addition to our understanding of the evolutionary particle architecture within the virosphere, expanding our knowledge of viral diversity and evolution.

## Materials and Methods

### Fungal strains

Strains CYG1-2 and CJP3-1 of *Ps. camelliae-sinensis* were isolated. from tea leaves (*C. sinensis (L.) O. Kuntze*) collected in Hubei province, China. They were identified based on molecular and morphological analyses (加Ref.He Yunqiang, Li Yan, Song Yulin, Hu Xingming, Liang Jinbo, Shafik Karim, Ni Deding, Xu

Wenxing*. Amplicon sequencing reveals novel fungal species responsible for a controversial tea disease. Journal of Fungi 2022, 8, 782) and subsequently purified using the hyphal-tipping technique (16). Strain XA-3 of *B. dothidea* was isolated from apple branches (*Malus domestica* Borkh. cv. ‘Fuji’) collected in Shandong province, China and was previously identified as being infected by BdRV1 in our previous studies (17).

### Extraction of dsRNAs and enzymatic treatments

For dsRNA extraction, mycelial plugs were inoculated in cellophane membranes on PDA (20% diced potatoes, 2% glucose, and 1.5% agar) plates and incubated at 25°C in the dark for 4–5 days. The mycelia were then collected, ground to a fine powder in liquid nitrogen, and subjected to dsRNA extraction and protein elimination using a patented method developed in our lab (18). The dsRNA preparations were further processed to remove the remaining ribosomal RNAs. Initially, aliquots of 200 ng of total dsRNA were treated with 10 U S1 nuclease (Thermo Scientific) at 37 °C for 1 h. Subsequently, the purified RNAs were analyzed by 1.5% agarose gel electrophoresis and visualized by staining with ethidium bromide. Each dsRNA band was excised separately and purified using a gel extraction kit (Qiagen, USA). The purified dsRNA samples were dissolved in DEPC-treated water and stored at -70 °C until use.

### Cloning and sequencing

The cDNA sequences of genomic dsRNAs were determined as previously described (19). Part of sequences of the dsRNAs were determined by cloning and sequencing amplicons generated by RT-PCR, using the random primers 05RACE-3RT and 05RACE-3 (Table S1). The 5′- and 3′-terminal sequences of the dsRNAs were obtained via cloning and sequencing of the RT-PCR amplicons, employing a standard RNA ligase-mediated RACE protocol, which included the use of PC3-T7loop and PC2 (Table S1). The oligonucleotide primers for RACE were designed using sequence information obtained from the randomly primed amplicons (20). At least three independent clones of each fragment were sequenced in both directions by Tsingke (Beijing) Co., Ltd, China.

Sequence similarity searches were performed using National Center for Biotechnology Information (NCBI) databases with the BLAST program. Multiple alignments of nucleic and amino-acid sequences and phylogenetic tree construction were conducted using MEGA version 11 (21). ORFs were deduced using ORF finder available at https://www.ncbi.nlm.nih.gov/search/all/?term=ORFfinder. The functions of the deduced proteins were predicted with SMART (Simple Modular Architecture Research Tool) online platform accessible at http://smart.embl-heidelberg.de/smart/set_mode.cgi?NORMAL=1.

### Purification of virus particles from mycelia

A mycelial plug was grown at 25°C in darkness for 8 days in sterilized cellophane membranes placed on PDA. Following harvest, 40 g mycelia were ground to a fine powder using liquid nitrogen, and the resulting powder underwent extraction as previously described (18). The crude or purified virus particle preparations were negatively stained with 2% sodium phosphotungstate on carbon-coated 200-mesh copper grids and subsequently examined by TEM (H-7000FA; Hitachi). The widths and lengths of the particles were measured using ImageJ, with the width of each particle was determined based on at least five measurements along randomly selected particles.

### Analysis of the dsRNAs and proteins from the viral particles

Purified virus-like particles underwent viral dsRNA extraction and analysis as previously described (18). Briefly, 50 μl sucrose suspension was collected from each fraction after sucrose gradient centrifugation and treated with phenol/chloroform/isoamyl alcohol (25:24:1) (pH 5.2) to remove viral proteins. The nucleic acids were then precipitated with ethanol, dissolved in DEPC-treated water, and analyzed by polyacrylamide gel electrophoresis.

Proteins extracted from each sucrose fraction were analyzed by 12% SDS-PAGE with 25 mM Tris-glycine and 0.1% SDS. Following electrophoresis, the gels were stained with Coomassie brilliant blue R-250 (Bio-Safe CBB; Bio-Rad, USA). Subsequently, the protein bands on the gel were individually excised and subjected to PMF analysis by Sangon Biotech (Shanghai) Co., Ltd, China, as previously described (22).

### Polyclonal antibody production and ISEM examination

The P5 protein was obtained and purified from prokaryotic expression as previously described (23). Briefly, amplified ORF5 fragments were inserted into the expression vector pET-28a, resulting in the reconstructed vector, pET-28a-P5-His. Histidine (His)-tagged recombinant proteins were subsequently affinity purified from *Escherichia coli* BL21 and dialysis for 2 h in room temperature. To generate polyclonal antibodies (PAb-P5), 300 μg of P5 protein was injected four times into two 5-week-old female BALB/C mice, which were obtained from the Laboratory Animal Centre, Huazhong Agriculture University, Hubei Province. The injection protocol followed previously established procedures (24). Protein preparations from both PcsPmV1-infected and - free isolates were extracted from sucrose fractions after sucrose gradient centrifugation, employing methods described in prior studies (18). These protein samples underwent indirect ELISA, as previously detailed, utilizing PAb-P5 dilutions ranging from 1:50 to 1:128,000 to determine the optimal titer (9). Western blotting and ISEM analysis were performed as previously described (25), utilizing PAb-P5 at dilutions ranging from 200- to 8000-fold.

### Immuno-gold labeling

Immuno-gold labeling was performed as previously described (26) with some modifications. Briefly, carbon-coated 200-mesh copper grids were floated on drops of the virus suspension samples for 3 min. The grids were treated with the primary antibody (PAb-P5) and allowed to incubate for 3 min. Subsequently, the grids were exposed to the second, which was incubated with the grid for 3 min (1/3 to 1/20 dilution of goat anti-rabbit 15 nm colloidal gold, Beijing Biosynthesis Biotech, Beijing, China). Finally, the grids were negatively stained and observed as described above.

### Elimination of PcsPmV1

Mycelial plugs of *Ps. camelliae-sinensis* strain CYG1-2 were cultured on PDA at 25°C under a 24 h photoperiod for more than 1 month until sporulation occurred. At the end of the culture, the conidia were collected and cultured on PDA plates at 25°C in darkness for 24 h. The resulting small colonies were individually transferred to new PDA plates for dsRNA extraction. The extracted dsRNA was analyzed by 1.5% agarose gel electrophoresis, and the bands were visualized by staining with ethidium bromide, and then further identified by RT-PCR. RT-PCR amplification was performed using a specific primer pair derived from the dsRNA 1 sequence (PcsPV1-1-1130F: 5′-CGACATCTCCCACTTCCTCC-3′; PcsPV1-1-1761R: 5′-CAGTCTCCTTCACCTTCAGC-3′), which generated a 632-bp fragment, using a PCR Thermal Cycler (Model PTC-100, MJ Research, USA) with an annealing temperature of 58°C.

## Ethics statement

This work was approved by the Research Ethics Committee, Huazhong Agricultural University, Hubei, China (HZAUMO-2024-0045), and carried out in accordance with the recommendations in the Guide for the Care and Use of Laboratory Animals from this committee.

## Data analysis

Descriptive statistics were determined, and X^2^-tests, one-way analysis of variance, and Tukey post hoc tests were performed using SPSS Statistics 17.0. p < 0.05 was considered to indicate statistical significance.

## Data availability

Sequence data supporting the findings of this study have been deposited in GenBank under accession numbers PP3594105-PP359410 for dsRNAs 1-6 of PcsPmV1, respectively. The remaining data are available within the article and its Supplementary Information files and from the corresponding author upon request.

## Acknowledgements

This work was supported by grants from Key R&D Program of Hubei (No. 2023BBB098) and the National Natural Science Foundation of China (32172475) to Wenxing Xu.

## Author Contributions

W.X. conceived this study, designed the investigation, and supervised the project. Z.H. conducted most of the experiments and wrote the manuscript, J.J. purified BdRV1, and checked its virions, and examined the related proteins. W.X. improved English, presentation, and discussion of the manuscript.

---

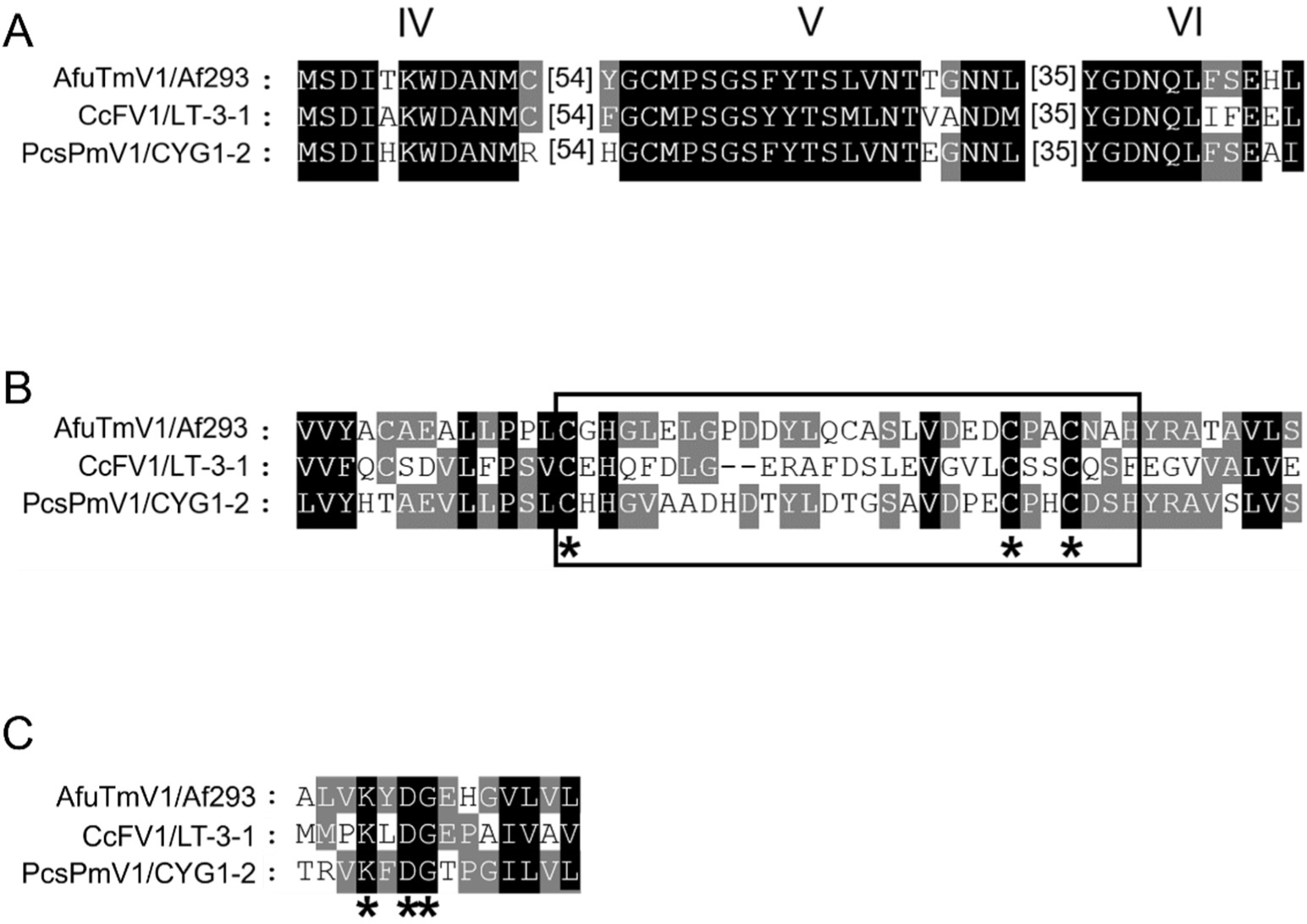

